# Circadian phase and sex shape swimming exercise responses and stereotyped behaviors in mice

**DOI:** 10.64898/2026.03.04.709589

**Authors:** Mónica D. Cortés Soto, Caoyuanhui Wang, Erin Kang, Sophia Martinez, Justin M. Toller, Hailey E. Vasquez, Samantha V. Herrera, Karina Alviña

## Abstract

Exercise provides broad health benefits, including improved emotional well-being and cognitive function. Emerging evidence suggests that exercising at different times during the day can have differential effects. However, how circadian phase and sex influence behavioral and physiological responses to exercise remains unclear. To address this question, we examined male and female wild-type mice maintained in either regular (REG, lights on/off at 7AM/7PM) or inverted (INV, lights off/on at 10AM/10PM) light cycles. Mice were then subjected to daily 20-min group swimming exercise sessions at ZT2-3 for 3 weeks. Exercised and sedentary controls mice were then subjected to an open field test (OFT) and blood corticosterone (CORT) measurements 24 hours post-exercise.

We quantified several behaviors during swimming: escape attempts, floating, climbing and collisions. We also identified a novel swimming behavior: floating with only nostrils-above-water events (NAWEs). We found that expression of these behaviors was differentially modulated by sex, light-cycle and their interaction. Notably, behavioral differences were more pronounced in REG mice (rest phase). REG mice also lost weight after exercise and had elevated CORT levels compared to mice kept in INV conditions (active phase). Interestingly, OFT behaviors showed significant differences primarily in INV mice, particularly females, when comparing exercised vs sedentary groups.

Our novel findings reveal that circadian rhythms and sex significantly interact to shape swimming exercise and stereotyped behaviors in mice. This emphasizes the need to consider the animals’ circadian phase when designing preclinical studies to match intended behavioral and physiological outcomes.

**HIGHLIGHTS:** Circadian phase and sex jointly shape swimming behavior patterns.

Newly identified swimming behavior is more prevalent during rest-phase

Rest⍰phase exercise produced stronger behavioral and physiological effects.

Rest-phase exercise resulted in weight loss and elevated stress markers.

Active-phase exercised females showed the strongest open field behavioral differences.

## INTRODUCTION

Physical exercise has been consistently linked to a wide range of health benefits including improving cardiovascular health, reducing risk of over 35 chronic conditions (among them type 2 diabetes and hypertension), and strengthening the immune system^1^. Beyond physical outcomes, exercise also has a positive impact on mental health. Studies demonstrate that people who exercise regularly show less symptoms of depression and anxiety and improve their cognitive function and executive control^2,3^. Furthermore, physical activity has been associated with enhanced neuroplasticity and cognitive health outcomes such as increased hippocampal neurogenesis^4^ and hippocampal volume concurrent with better memory performance^5^. These findings support the notion that exercise is a key protective factor to be considered across the lifespan, with important considerations for cognitive aging and neurodegenerative diseases^6^. This evidence suggests that exercise can be used as an effective non-pharmacological and accessible tool to promote health and disease prevention in a holistic manner.

Emerging evidence has highlighted the importance of exercise timing. For example, a recent meta-analysis concluded that a single session of aerobic exercise in the evening promoted a greater reduction in blood pressure and cardiac workload post-exercise compared to exercising in the morning^7^. Another meta-analysis explored the effects of time of day on dynamic short-duration maximal exercise performance (DSDME) and found that it peaked during the afternoon, between 4-8PM^8^. However, other studies have shown the opposite effect. For instance, high intensity-interval training (HIIT) exercise was more effective in improving cognitive function when executed in the morning compared to the afternoon^9^. Other studies have shown no significant effect of time of day on cognitive function^10^. While these discrepancies might be related to differences in the type of exercise evaluated and/or the population included (different age, gender, etc.), these results (and many others) nevertheless provide evidence that circadian rhythm influences the potential benefits of exercise.

Circadian rhythms control the timing of physiological, biochemical, and behavioral processes in a 24-hour periodicity^11^. At a molecular level, circadian rhythms consist of a regulatory network of transcriptional-translational feedback loops. The core activators, CLOCK (Circadian locomotor output control kaput) and BMAL1 (Brain muscle arnt-like1), form heterodimers that bind to the promoters of PER1/2 (Period) and CRY1/2 (Cryptochrome) genes, initiating their transcription. Once translated, PER and CRY proteins accumulate in the cytoplasm, then re-enter the nucleus where they inhibit CLOCK–BMAL1 activity, thereby repressing their own transcription and generating oscillations of approximately 24 hours^12^. This molecular clock is present in virtually all cells in the body. Additionally, there is a central clock located in the suprachiasmatic nucleus (SCN) of the hypothalamus, which receives direct photic input from the retina, allowing environmental light cues to reset and synchronize the internal clock with the day-night cycle^13^. Therefore, the most important cue, also known as Zeitgeber (ZT), for the entrainment of the central clock, is light^14^. However, non-photic cues, such as feeding, temperature, and exercise, also play essential roles in the circadian rhythms entrainment^15^, particularly in peripheral organs and tissues such as skeletal muscle^16^. The skeletal muscle clock can be entrained indirectly by light input (via the central clock) and directly by feeding and activity^16,17^. Moreover, when these rhythms are disrupted, skeletal muscles exhibit adverse cellular changes that lead to reduced muscle performance^18^. More recent studies have shown that exercise can act as a time-cue to entrain the skeletal muscle clock. Mice that ran during the inactive phase of the light cycle, either voluntarily or forced, showed a significant phase shift in PER2 gene expression^19,20^. These findings further support the notion that exercise could function as a potential chronotherapeutic intervention^21^.

The main objective of this study was to investigate how the timing of exercise during the light cycle influences behavioral and physiological responses in mice. We used both male and female wild-type mice kept in opposite light cycles and compared exercise performance during daily swimming sessions. After exercise, we monitored exploratory behaviors in the open field test and assessed baseline corticosterone levels. Our results showed that sex and circadian phase jointly shape swimming performance, including newly identified and stereotyped behaviors.

## METHODS

### Animals

All procedures detailed here were approved by the University of Florida Institutional Animal Care and Use Committee (IACUC), Animal Protocol # IACUC202300000538, and follow the U.S. National Institutes of Health (NIH) guidelines for animal research.

Young adult wild-type mice (8-10 weeks old, C57BL/6J background) were obtained from our own breeding colony. Mice were always group-housed (2 – 5 mice per cage) in ventilated cages inside air-conditioned rooms kept at 21-23°C (70-74°F) and 40-60% humidity. Two weeks before experiments started, mice were transferred to specific rooms with different 12h/12h light cycles and kept unperturbed until the beginning of the experiments. For the regular daylight room (referred to as REG), the lights were automatically switched ON at 7:00 AM and OFF at 7:00 PM. For the inverted light-cycle room that was kept in dark conditions during the day (referred to as INV), lights were turned OFF at 10:00 AM and back ON at 10:00 PM. Throughout the experiment, mice had ad libitum access to standard laboratory food and water. All behavioral experiments were performed in a dimly lit separate room inside the animal facility, in proximity to the room where mice were kept.

### Swimming exercise protocol

Mice were subjected to daily swimming exercise in groups of cage mates. We used a large square shaped plastic tank measuring 45.7cm per side (∼18in, total area ∼324 in^2^) with a plastic divider placed diagonally in the middle to divide it into two separate triangle-shaped areas. The tank was filled with water to a height of 5-6in, ensuring that mice could not stand on their feet. The water temperature was kept at 32°C. To reduce surface tension (and encourage active swimming) we added a drop of liquid soap to the water before starting each session.

The swimming protocol included two phases: habituation and training. During the habituation period, mice were subjected to swimming for 5 minutes the first day, then adding 5 min each consecutive day until completing 20 minutes of total swimming (day 4). After the 4th day, mice swam for 20 min daily for the next consecutive 21 days. At the end of each swimming session, mice were gently removed from the water and placed on a clean cage covered with paper towels kept on top of a heat pad. Exercise was performed at the same time every day, starting at ZT2-3 for both REG mice (10AM – 12PM) and INV mice (12–2 PM). Mice were transported to the behavior room 30 min before the beginning of their swimming session every day. During this time all mice were weighed (daily), including the sedentary age- and sex-matched controls that were then returned to the room where they were kept.

While swimming, mice were always monitored and their different behaviors manually tallied. Six key swimming behaviors were identified and quantified (Table 1). These included: escape attempts, floating (defined as >50% of the body above water), climbing, collisions, defecation, and floating while body was submerged (i.e., nostrils-above-water events or NAWEs), defined as follows:

**Table 1.** Classification and definition of specific behaviors documented during swimming sessions.

| Behavior | Classification | Definition |
| --- | --- | --- |
| Escape attempts | Active | Movement directed at grasping the wall, divider or corners of the water tank, trying to stop swimming for >3 s. |
| Collisions | Active | Two or more mice bump into each other while actively swimming or a mouse step onto another while their bodies touch during active swimming. |
| Climbing | Active | Mouse flaps its front and hind limbs pushing itself up above water. This behavior was usually accompanied by submersion and resurfacing and then active swimming. |
| Floating | Passive | Mice exhibiting minimal or no active movement for >3 s while >50% of its body is above water, in a mostly horizontal position. |
| Nostrils above water events (NAWEs) | Passive | Mouse floating while keeping only its nostrils above the water (no active swimming) and its body is almost vertical under water. |
| Defecation | Passive | After finishing each group swimming session, the total number of fecal particles was counted and divided by the number of mice to calculate defecation per mouse. Based on our observations, 1-3 fecal particles per mouse were considered a low defecation range; 4-6 particles were considered a medium defecation range; and >7 particles were classified as high. |

### Open field test

The day after the last day of swimming, mice were subjected to an open field test (OFT), scheduled at the same time as the mice usually exercised. Sedentary and exercised mice were brought to the behavior room, weighed and left unperturbed for 30 min before the test. For the OFT, mice were placed individually in the center of a square arena consisting of a plastic container of 45.7 cm per side (∼18in) and allowed to fully explore for 10 minutes while videotaping for further behavioral analysis. Data acquisition and analysis was conducted using Anymaze Software (Version 7.4; Stoelting Co., Wood Dale, IL).

### Tissue Collection

Mice were anesthetized with 3% isoflurane, USP (NDC 46066-115-03, PIVETAL) and decapitated immediately after behavioral tests. Trunk blood samples were collected and centrifuged for 10 min at 2000X g, then serum was retrieved and stored at –80°F until use.

### Corticosterone ELISA Assay

Serum corticosterone was measured in duplicate using a commercial chemiluminescent immunoassay kit (Cat #K014-C, Arbor Assays, Ann Arbor, MI, USA). Diluted serum samples were processed according to the manufacturer’s instructions. For serum sample dilution, 5 μL of serum was incubated with 5 μL of dissociation reagent, and diluted 250X with assay buffer (500X dilution, reagent provided by commercial kits).

### Data analysis

All statistical analyses were performed using Origin software (OriginLab Corporation). Unless otherwise noted, all data are presented as Mean ± SEM. Depending on the number of factors included, Two- or Three-way ANOVA was conducted, followed by Tukey’s post-hoc test for means comparison. P≤0.05 was considered statistically significant.

## RESULTS

### Exercise induces body weight loss or reduced weight gain in mice maintained in REG cycle

To monitor overall health during the 21-day swimming protocol, we documented body weight (BW) for each mouse before every daily swimming session. We also weighed mice in the sedentary control groups at the same time, then returning them to their home cages and leaving them unperturbed afterwards. Figures 1A and B show the average daily BW per group during the swimming protocol separated by light cycle. We normalized BW values to each mouse’s starting weight to account for baseline sex-dependent differences. Our results showed that exercise led to a significant reduction in BW gain in mice kept in REG light compared to their sedentary (SED) counterparts (Figure 1A), whereas mice kept in INV conditions (Figure 1B) either showed no change (females) or their BW increased after exercise (males). Furthermore, we observed clear ∼5day fluctuations in the BW of female mice kept in REG conditions that were not apparent in female mice kept in INV light cycle conditions, suggesting a pattern likely related to the duration of the estrous cycle^23^. Statistical analysis revealed that both sex and exercise had a strong individual effect (p<0.0001 for both, Three-way ANOVA). Moreover, interactions between all factors were significantly different: p=0.002 for light cycle*sex, p<0.0001 for light cycle*exercise and p<0.0001 for sex*exercise, while the three-way interaction between light cycle*sex*exercise was not significantly different (p=0.717).

**Figure 1.**
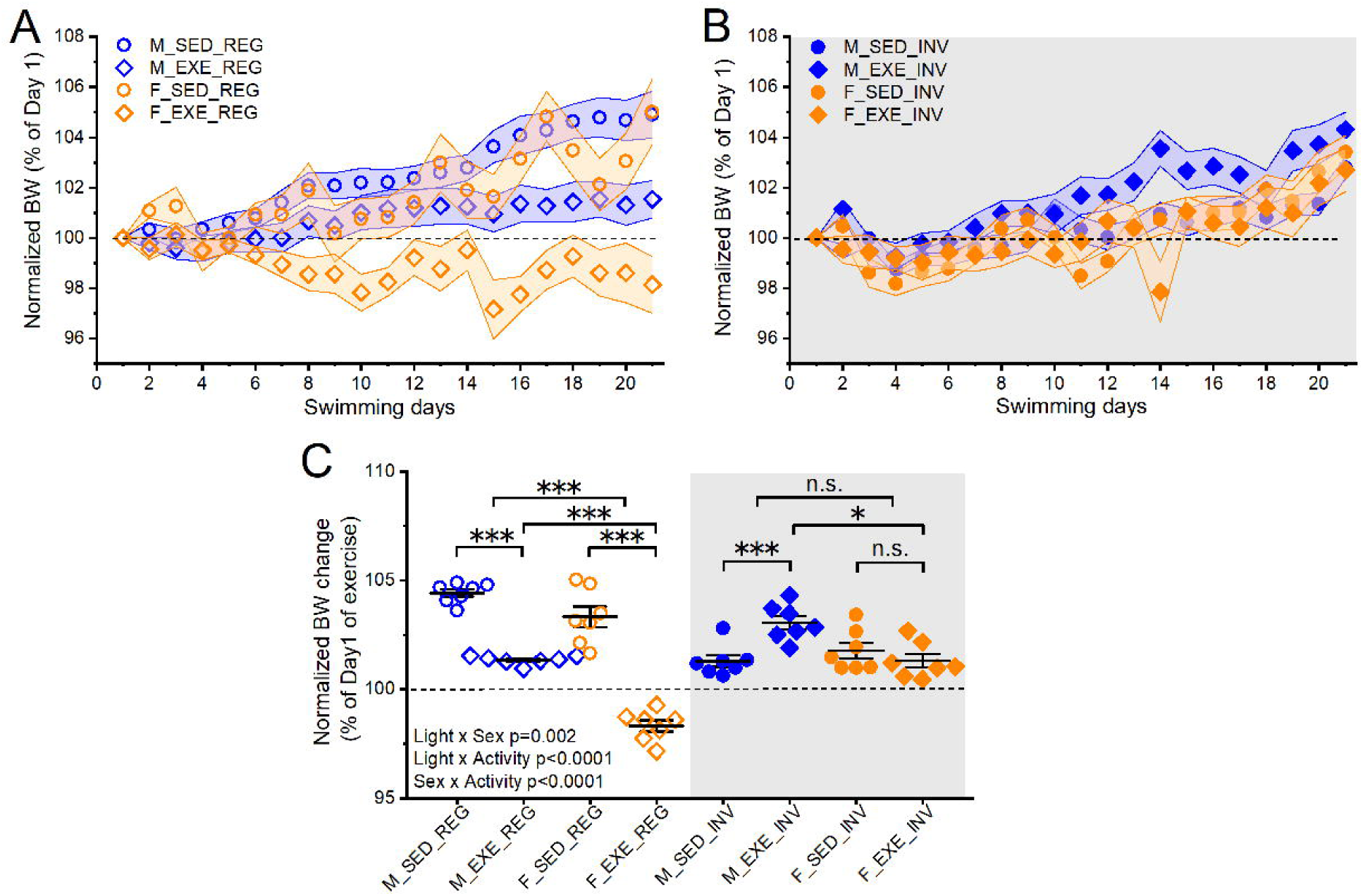
Normalized body weight (BW) for sedentary (SED) or exercised (EXE) male (blue symbols) and female (orange symbols) mice kept in the regular (A, open symbols) or inverted (B, closed symbols) light cycle over the 21 days of the swimming protocol. Each data point represents the group mean and the shaded area around each data point depicts ±S.E.M. The last 7 days of swimming were averaged and plotted per group (C). Statistical analysis was conducted by Three-way ANOVA and Tukey’s test post hoc for means comparison. (***) denotes statistical significance with p<0.001, (*) denotes p<0.05, n.s. denotes “not significant” (p>0.05).

We then averaged BW changes over the last 7 days of exercise per group. As shown in Figure 1C, sedentary males in REG conditions (M-SED-REG, n=15 mice) gained significantly more weight than exercised males (M-EXE-REG, n=17 mice). Sedentary females in the same light cycle (F-SED-REG, n=13 mice) gained almost as much weight as the sedentary males, whereas exercised females (F-EXE-REG, n=16 mice) were the only group that lost a significant amount of weight during the swimming protocol (average BW change in M-SED-REG = 104.4±0.2%; M-EXE-REG = 101.4±0.1%; F-SED-REG= 103.3±0.5% and F-EXE-REG = 98.3±0.3%; p<0.001, Three-way ANOVA, Tukey’s Test for means comparison). In contrast, sedentary male mice kept in NIGHT conditions (M-SED-NIGHT, n=16 mice) did not gain as much weight as M-SED-REG, while exercised INV males gained weight after exercise (M-EXE-INV, n=17 mice). The average BW change in M-SED-INV and M-EXE-INV was 101.3±0.3% vs 103.1±0.3% respectively (p<0.001). Furthermore, exercise had no effect on BW in female mice kept in INV light conditions as the average BW change in F-SED-INV was 101.8±0.4% (n=14 mice) and in F-EXE-INV it was 101.3±0.3% (n=12 mice, p=0.933 between SED and EXE female INV groups).

Overall, our results demonstrated a significant modulation of BW after swimming exercise, and second, this change was strongly influenced by the interaction between light cycle and sex. In fact, male mice exhibited the most pronounced effect as males kept in REG showed completely opposite responses to exercise compared to male mice in INV conditions.

### Swimming behavior is modulated by the interaction between light cycle and sex

The changes in BW previously described might reflect metabolic differences that could have important effects on baseline and induced behaviors. Therefore, to accurately characterize potential differences in behavior during swimming sessions, we established specific parameters based on careful observations by multiple researchers in our team (as described in Methods). From this analysis, we developed our own swimming ethogram consisting of 6 reliably distinctive and identifiable behaviors: escape attempts, floating (defined as keeping >50% of the body above water while floating for ≥3s), climbing events, collisions, defecation, and nostrils-above-water events (NAWEs). These swimming behaviors were manually scored and tallied during the last 10 days of the swimming protocol to reduce the effect of habituation/novelty expected during the first few days the swimming protocol. Additionally, the scoring process was conducted during the last 15 min of each 20 min session (to also reduce the influence of the first minutes of acute response to the water). Because mice swam in groups of cage mates, and numbers of mice per cage differed (ranging from 2 to 5), we divided the total number of counted events by the number of mice in each cage. Behaviors were further categorized into active or passive, with escape attempts, collisions, and climbing being considered active swimming behaviors (Figure 2). Notably, because the swimming sessions were not videotaped, we did not include in this analysis the time that mice spent actively swimming, though it can be inferred from the combined scored behaviors.

**Figure 2.**
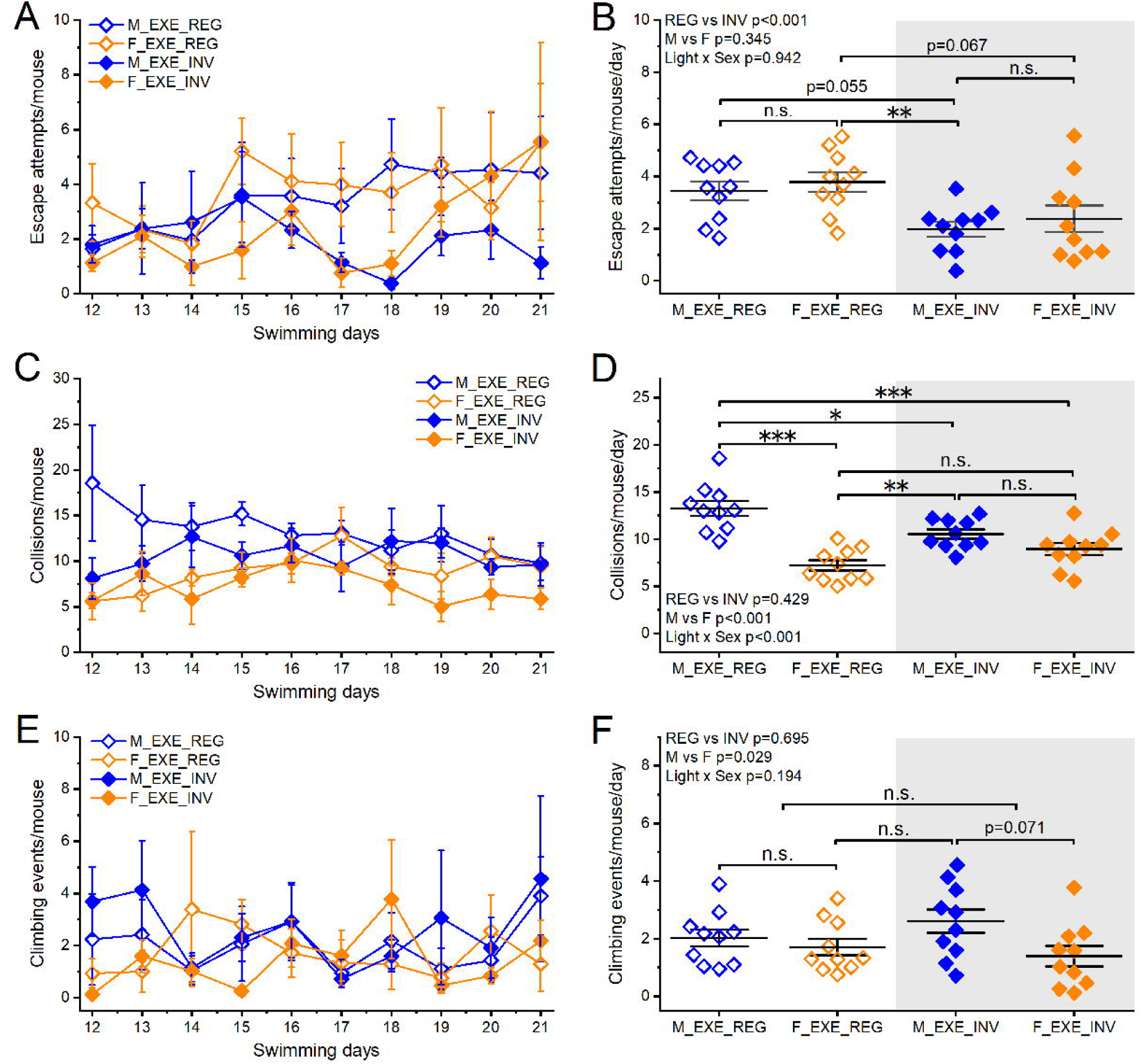
Active swimming behaviors over the span of 10 days of daily swimming in mice kept in regular (REG) or inverted (INV) light cycle. A and B show the daily average per group for escape attempts and the mean ±S.E.M. for the last 7 days, respectively. C and D show the daily average per group for collision and the mean ±S.E.M. for the last 7 days, respectively. E and F show the daily average per group for climbing attempts and the mean ±S.E.M. for the last 7 days, respectively. Statistical analysis was conducted by Three-way ANOVA and Tukey’s test post hoc for means comparison. (***) denotes statistical significance with p<0.001, (**) denotes p<0.01 and “n.s.” denotes “not significant” (p>0.05).

Daily averages of escape attempts per group are depicted in Figure 2A, which shows that mice kept in REG conditions displayed significantly more escape attempts compared to INV groups, regardless of sex. Figure 2B shows the average number of events per mouse over the 10 days of swimming plotted in 2A, for each group. On average, M- and F-EXE-REG made more escape attempts (3.4±0.4 and 3.8±0.4 events per mice per day, respectively, p=0.923 between M and F groups, Two-way ANOVA and Tukey’s Test for means comparison) compared to mice in the M-and F-EXE-INV groups (1.97±0.3 and 2.4±0.5 events per mice per day, respectively, p=0.885 between M and F groups). While the difference in escape attempts between sexes within each light cycle did not reach statistical significance (p=0.345, overall Two-way ANOVA), comparing between sexes across light cycles there was a significant reduction in escape attempts in mice kept in INV conditions (p<0.001 between REG and INV groups, overall Two-way ANOVA). Similarly, the interaction between sex and light cycle was not statistically significant (p=0.942).

In contrast, we observed a significant sex dependent effect (p<0.001; overall Two-way ANOVA) in the number of collision events with no effect of light cycle (p=0.429, Figure 2C-D). On average, M-EXE-REG engaged in more collisions (13.3±0.8 events per mice per day) than F-EXE-REG (7.2±0.6 events per mice per day, p<0.001). This difference was not exhibited in mice kept in INV conditions, as the number of collisions did not differ between M- and F-EXE-INV (10.5±0.5 vs. 8.9±0.7 events per mice per day, respectively, p=0.43). Moreover, the interaction between sex and light cycle was highly statistically significant for collisions (p<0.001). Our analysis also showed a comparable effect on climbing events (Figure 2E-F). This behavior was significantly influenced by sex (p=0.029, overall Two-way ANOVA) while light cycle had no individual effect (p=0.695). On average, M- and F-EXE-REG showed similar levels of climbing behavior (2.02±0.3 vs. 1.7±0.3 events per mice per day, p=0.91), while male mice in INV conditions showed more climbing than F-EXE-INV (2.6±0.4 vs 1.4±0.4 events per mice per day), however, this difference did not reach statistical significance (p=0.071, Two-way ANOVA and Tukey’s Test for means comparison).

We also identified behaviors during swimming that we considered passive, such as floating, NAWEs and defecation. In this case, we observed a significant sex dependent effect (p=0.046) in the number of floating events with no effect of light cycle (p=0.764, overall Two-way ANOVA). Figures 3A and B show that the most pronounced difference was observed between M- and F kept in REG light cycle. The average number of floating events by the M-EXE-REG group was 5.7±1.2 events per mice per day, whereas the F-EXE-REG group more than doubled that number (12.9±0.9 events per mice per day, p=0.0054, Two-way ANOVA and Tukey’s Test for means comparison). In contrast, this sex dependent difference was abolished in mice kept in INV conditions, as the number of floating events between M- and F-EXE-INV was 10.4±1.9 vs. 9.1±1.4 events per mice per day, respectively (p=0.916). Moreover, the interaction between sex and light cycle was highly statistically significant for floating events (p=0.005, overall Two-way ANOVA).

**Figure 3.**
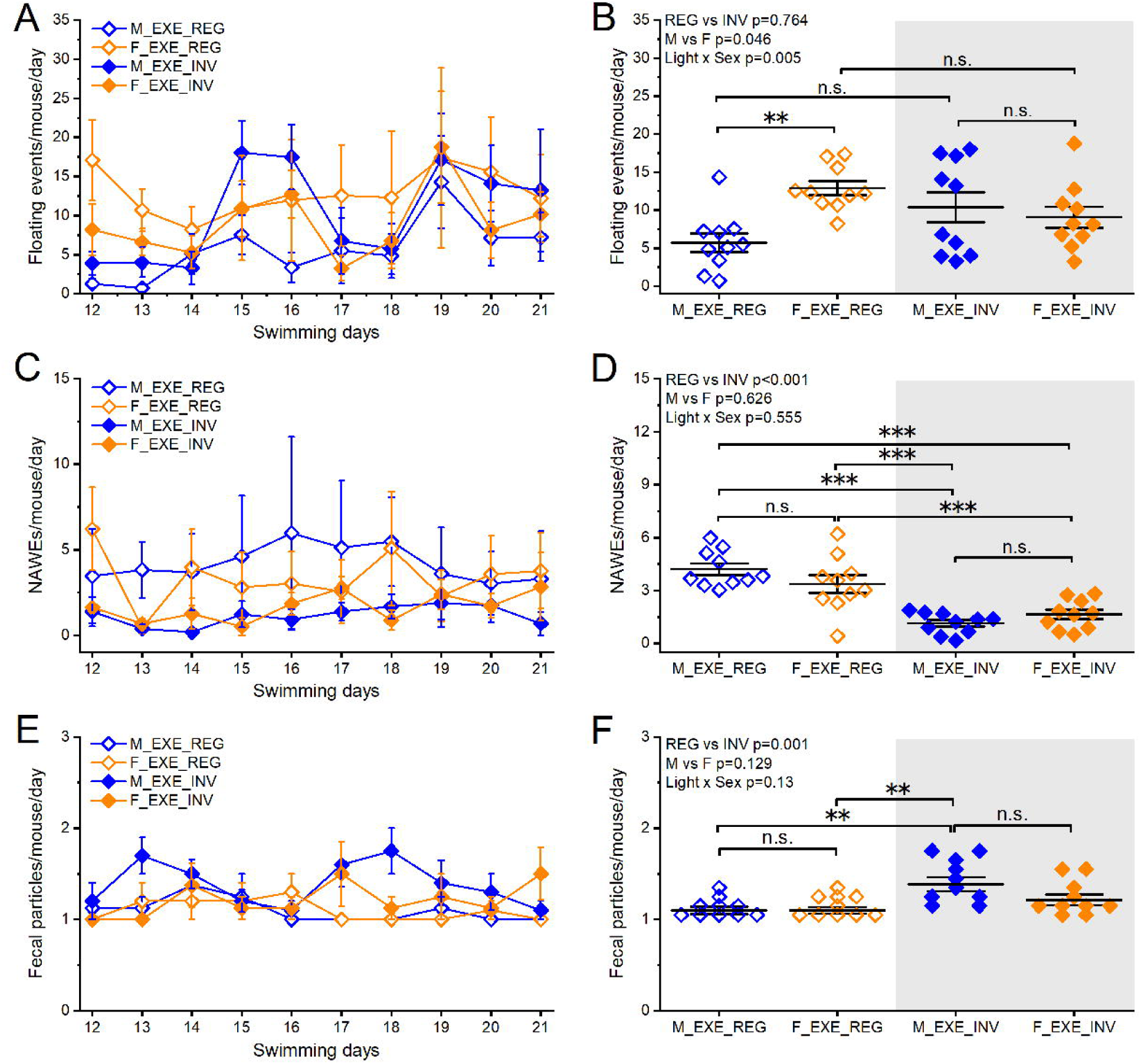
Passive swimming behaviors over the span of 10 days of daily swimming in mice kept in regular (REG) or inverted (INV) light cycle. A and B show the daily average per group for floating events and the mean ±S.E.M. for the last 7 days, respectively. C and D show the daily average per group for nostrils above water events (NAWEs) and the mean ±S.E.M. for the last 7 days, respectively. E and F show the daily average per group for fecal particles and the mean ±S.E.M. for the last 7 days, respectively. Statistical analysis was conducted by Three-way ANOVA and Tukey’s test post hoc for means comparison. (***) denotes statistical significance with p<0.001, (**) denotes p<0.01 and “n.s.” denotes “not significant” (p>0.05).

Our behavioral observations identified a highly stereotyped form of floating behavior that most mice displayed at some point. We defined this behavior as “nostrils above water events” or NAWE as this was the exact position that mice adopted, i.e., their body was almost vertical and fully submerged except for the nostrils. Our quantification showed a clear effect of light cycle as mice in REG conditions (both males and females) displayed significantly more NAWEs than mice in the inverted light cycle (p<0.0001, overall Two-way ANOVA). Conversely, sex had no significant effect (p=0.626), and there was no significant interaction between light cycle and sex (p=0.555). As shown in Figure 3C-D, the number of NAWEs displayed by M-EXE-REGs was 4.2±0.3 events per mice per day, whereas the F-EXE-REG group showed 3.4±0.9 events per mice per day (p=0.315, Two-way ANOVA and Tukey’s Test for means comparison). Similarly, no differences were observed in the number of NAWEs between M- and F-EXE-INV (1.2±0.2 vs. 1.6±0.3 NAWEs per mice per day, respectively, p=0.72).

Lastly, at the end of each swimming session, we quantified the number of fecal particles left in the water tank, as an additional readout potentially related to stress coping behaviors (Figure 3E-F). In this case, there was a statistically significant effect of light cycle as mice kept in REG produced fewer fecal particles than mice kept in INV conditions (p=0.001), regardless of sex (p=0.129), with no statistically significant interaction between these two factors (p=0.13, overall Two-way ANOVA. On average, the number of particles per mouse for the M- and F-EXE-REG groups was 1.1±0.04 and 1.1±0.03 per mouse per day, respectively (p=0.999). Similarly, for M-and F-EXE-INV groups, the average was 1.4±0.08 and 1.2±0.06 particles per mouse per day, respectively (p=0.5).

In summary, our ethogram of swimming behaviors indicates that mice display a series of distinctive behaviors that are strongly modulated by sex or light cycle, and in some cases, by a significant interaction between these two factors.

### Coping mechanisms during swimming exercise

To further analyze the behaviors identified during swimming exercise, we calculated an “immobility score” by simply adding together the average values of floating events, NAWEs and escape attempts per day per group (i.e., all behaviors that resulted in reduced or no movement during swimming). Note that these scores are reported without S.E.M. value as they represent the sum of average values for each quantified behavior. As shown in Figure 4A, which depicts consecutive daily immobility scores during the last ten days of swimming, immobility scores slightly increased over time, suggesting that mice might have learned to conserve energy rather than to continue swimming as part of their coping strategies ^24^. Regardless of this change, the average per experimental group showed significant sex-dependent differences that were only observed in mice kept in REG conditions (Figure 4B). Importantly, although the overall effects of sex and light cycle did not reach statistical significance when analyzed independently (p=0.053 between REG and INV, and p=0.072 between M and F, overall Two-way ANOVA), there was a significant interaction between these two factors (p=0.044), suggesting that coping mechanisms are modulated by both sex and circadian influences. Specifically, the immobility score for M-EXE-REG mice was 13.4±1.5 whereas for the F-EXE-REG group it was 20.04±1.3 (p=0.041, Two-way ANOVA and Tukey’s Test for means comparison). On the contrary, no differences were observed in the immobility score for M- and F-EXE-INV groups (13.5±2.1 vs. 13.1±1.8, respectively, p=0.998).

**Figure 4.**
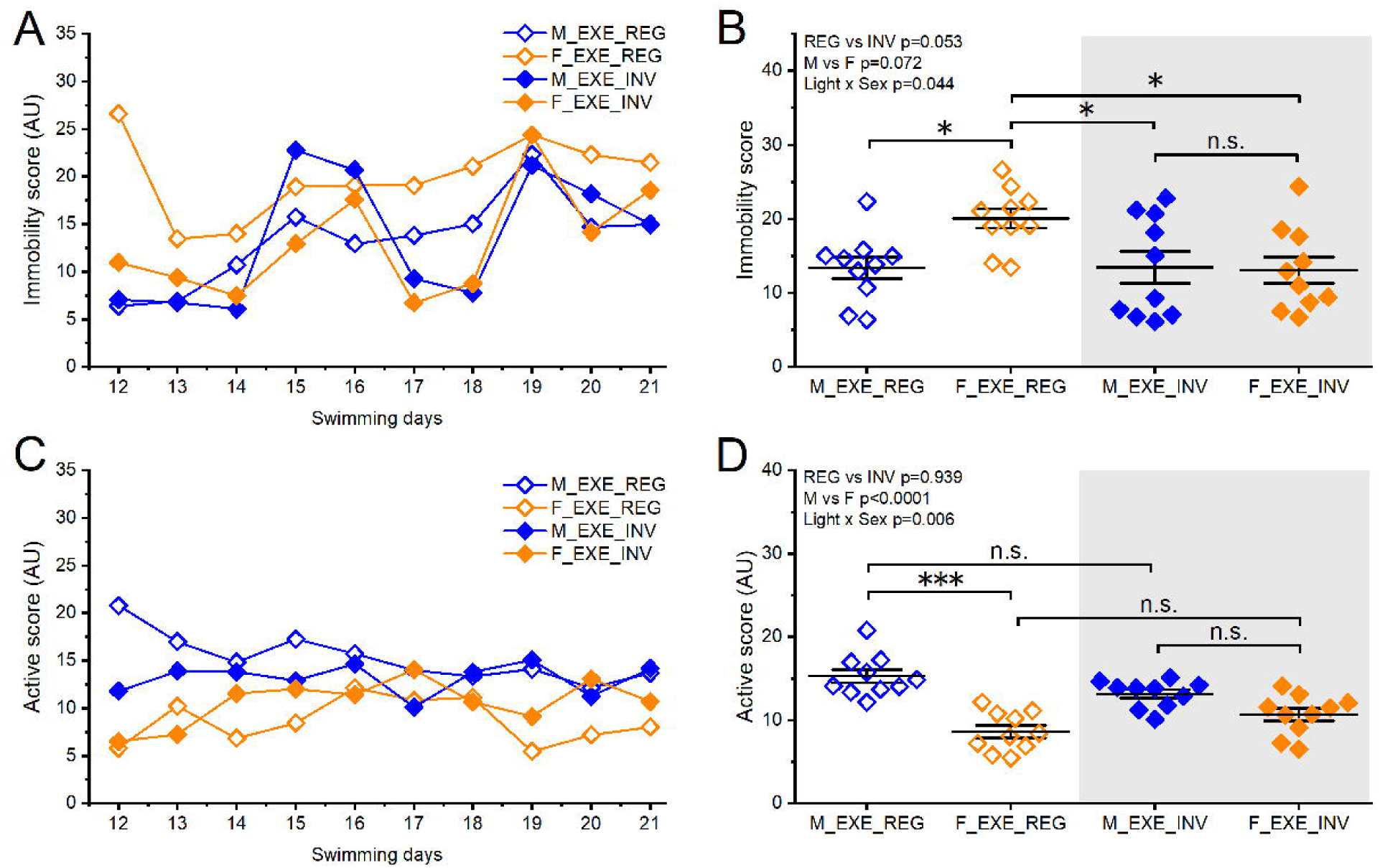
Daily immobility and mobility scores per group. A and B show daily averages for immobility score and mean ±S.E.M. for the last 7 days of swimming, respectively. C and D show daily averages for mobility score and mean ±S.E.M. for the last 7 days of swimming, respectively. Statistical analysis was conducted by Three-way ANOVA and Tukey’s test post hoc for means comparison. (***) denotes statistical significance with p<0.001, (*) denotes p<0.05 and “n.s.” denotes “not significant” (p>0.05).

To determine if active behaviors were equally modulated as passive, we calculated an “active score” by adding daily values of climbing events and collisions. As shown in Figure 4C, and in contrast to immobility scores, the active score showed no change over time as all groups displayed similar daily values throughout the last 10 days of swimming. However, averaging these values per experimental group revealed a similar trend to the immobility score, with significant differences observed only between M and F mice kept in REG conditions while M and F in INV conditions showed comparable active scores (Figure 4D). The average active score for M-EXE-REG mice was 15.2±0.8 compared with 8.6±0.7 for F-EXE-REG mice (p<0.0001, Two-way ANOVA and Tukey’s Test for means comparison). While there was a trend to lower active score for F-EXE-INV compared to M-EXE-INV, this difference was not statistically different (active score for M-EXE-INV was 13.1±0.5 vs. 10.7±0.8 for F-EXE-INV, p=0.08). Furthermore, compared to immobility score, our statistical analysis also revealed a strong interaction between sex and light cycle (p=0.006, overall Two-way ANOVA).

Taken together, these findings highlight the significant interaction between sex and light cycle in determining swimming behavior and potential coping mechanisms. This was particularly evident in the increasing immobility score as more swimming sessions were conducted.

### Exploratory behaviors in the open field test

To investigate the combined effects of light cycle, sex, and exercise on baseline locomotion and exploratory behaviors, we conducted an open field test (OFT) 24h after the last day of swimming (Figure 5), comparing between exercised mice and sedentary controls. To avoid additional daylight influence, we performed the OFT at the same time of the day that swimming sessions were conducted during the swimming protocol (at ZT 2-3). Our analysis showed that only exercise had a significant influence on the total distance traveled in the OFT when evaluated independently (p=0.026, overall Three-way ANOVA), while sex had no effect (p=0.322) and light cycle did not reach statistical level on its own but was very close (p=0.066). Furthermore, light cycle and exercise had a significant interaction (p=0.017). When comparing experimental groups (Figure 5A), the only group that showed a significant effect of exercise was the female mice kept in INV conditions, which moved more than the SED controls kept in the same light cycle. On average, F-SED-INV covered 30.4±1.5m vs. 40.6±1.8m for the F-EXE-INV group (p=0.038, Three-way ANOVA and Tukey’s Test for means comparison).

**Figure 5.**
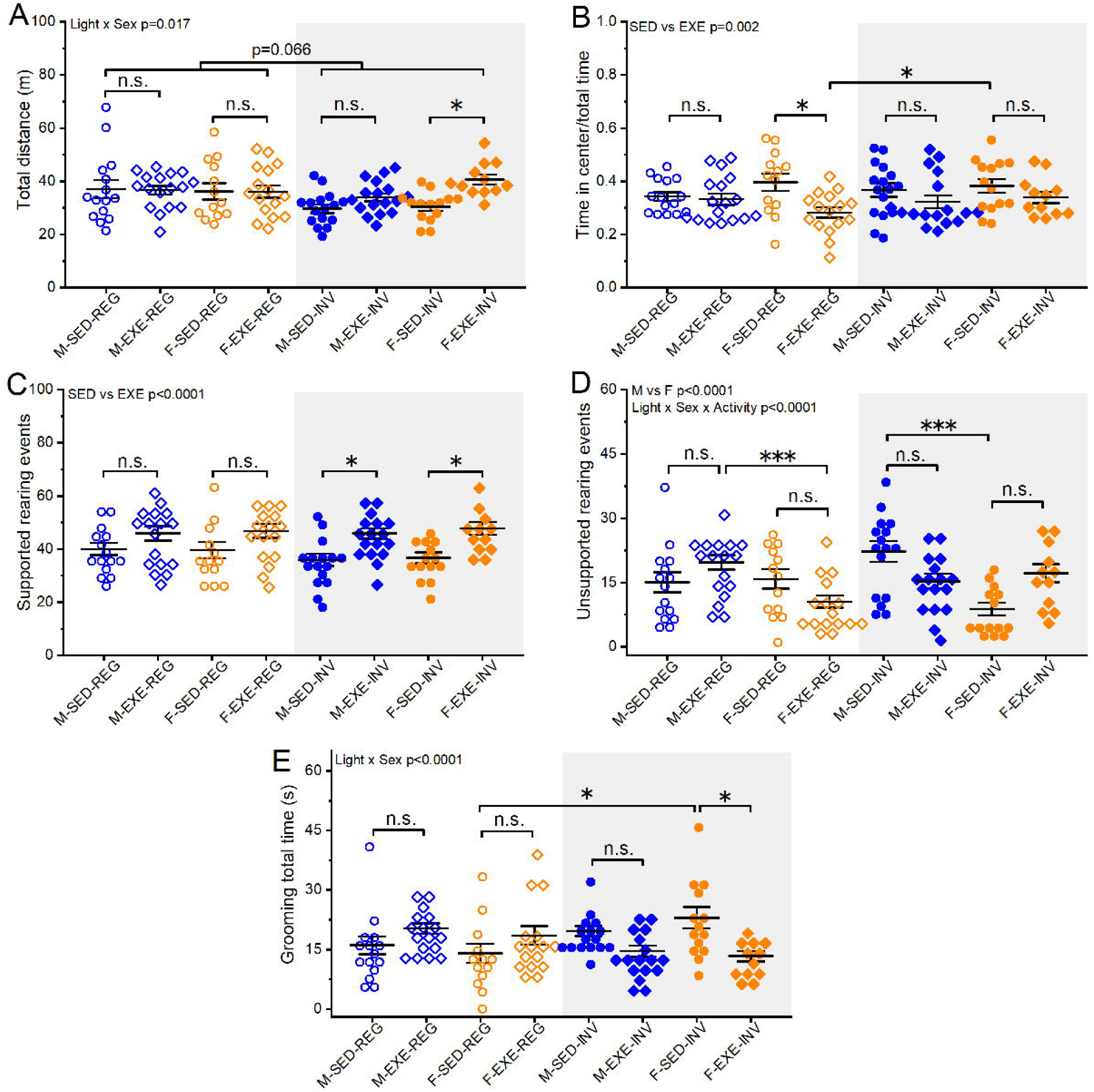
Analysis of locomotor and exploratory behaviors in the open field test (OFT), including total movement (A), time spent in the center (B), supported and unsupported rearing events (C and D) and grooming time (E). Statistical analysis was conducted by Three-way ANOVA and Tukey’s test post hoc for means comparison. (***) denotes statistical significance with p<0.001, (*) denotes p<0.05 and “n.s.” denotes “not significant” (p>0.05).

We then quantified the time spent in the center of the OFT as a well-established proxy for anxiety-like behaviors in rodents^25^. Figure 5B shows that exercise was the only independent factor that significantly impacted the ratio between time spent in the center vs total time in the OF (p=0.002, overall Three-way ANOVA). Neither light cycle nor sex had a comparable individual effect (p=0.394 and 0.584, respectively) or by interaction (light cycle*sex p=0.662; light cycle*exercise p=0.583; sex*exercise p=0.128; light cycle*sex*exercise p=0.12). Furthermore, when comparing all groups, we only observed a significant effect in females kept in REG conditions, which spent less time in the center of the OF, suggesting an anxiogenic effect of exercise that was abolished in mice kept in INV conditions. On average, F-SED-REG spent close to 40% (0.4±0.03) of their time in the center vs. 28% (0.28±0.02) for the F-EXE-REG group (p=0.016, Three-way ANOVA and Tukey’s Test for means comparison).

During the OFT we also quantified other well-established exploratory behaviors such as rearing and grooming. While sex-dependent effects were observed, light cycle did not impact any of these behaviors when analyzed separately. Rearing behavior was quantified as either supported (i.e., leaning against the OF wall) or unsupported (i.e., standing on hindlimbs with no support). This distinction showed important differences, as only supported rearing was strongly modulated by exercise (p<0.0001, overall Three-way ANOVA, Figure 5C), whereas unsupported rearing was not affected (p=0.88, Figure 5D). When analyzing group differences in supported rearing, while all exercised mice displayed more rearing events, only the ones kept in INV conditions (both males and females) reached statistically significant differences. On average, mice in the M-SED-REG and M-EXE-REG groups showed 40.1±2.3 vs 45.9±2.7 supported rearing events (p=0.591, Three-way ANOVA and Tukey’s Test for means comparison), while F-SED-REG and F-EXE-REG mice showed 39.6±3.1 vs 46.9±2.6 supported rearing events (p=0.389). Similarly, mice in the M-SED-INV and M-EXE-INV groups showed 35.9±2.3 vs 45.9±1.9 supported rearing events (p=0.043), while F-SED-INV and F-EXE-INV mice showed 36.8±1.9 vs 47.8±2.4 supported rearing events (p=0.05).

Unsupported rearing events showed very different patterns compared to supported rearing. For instance, unsupported rearing showed a strong sex-dependent effect (p<0.0001, overall Three-way ANOVA, Figure 5D), while supported rearing did not (p=0.652, Figure 5C). In addition, light cycle had no major modulatory effect when analyzed separately (p=0.671), however the interaction between all three factors (light cycle*sex*exercise) was strongly significant (p<0.0001). The analysis of exercise-dependent effects showed no significant changes regardless of sex and light cycle, likely due to large variability within each group. On average, mice in the M-SED-REG and M-EXE-REG groups showed 15.1±2.3 vs 19.7±1.7 unsupported rearing events (p=0.634), while F-SED-REG and F-EXE-REG mice showed 15.8±2.7 vs10.6±1.5 unsupported rearing events (p=0.546). Similarly, mice in the M-SED-INV and M-EXE-INV groups showed 22.3±2.4 vs 15.3±1.7 unsupported rearing events (p=0.132), while F-SED-INV and F-EXE-INV mice showed 8.8±1.5 vs 17.2±2.2 unsupported rearing events (p=0.083).

Lastly, we quantified the total time grooming during the OFT. We focused on total time grooming instead of counting grooming events as these events can have different durations and thus the number of events might not accurately reflect the overall grooming behavior. As shown in Figure 5E, none of the factors we considered (light cycle*sex*exercise) had a significant effect individually (p=0.769 for light cycle, p=0.761 for sex, and p=0.271 for exercise; overall Three-way ANOVA). However, there was significant interaction between light cycle and exercise (p<0.0001). When we analyzed individual groups, the only significant difference was observed between female mice kept in INV conditions. On average, mice in the M-SED-REG and M-EXE-REG groups groomed for 16.04±2.3s vs 20.3±1.3s (p=0.713), while F-SED-REG and F-EXE-REG mice spent 14.1±2.4s and 18.5±2.3s grooming, respectively (p=0.715). Similarly, mice in the M-SED-INV and M-EXE-INV groups groomed for 19.7±1.3s and 14.6±1.4s (p=0.472), while F-SED-INV and F-EXE-INV mice spent 22.9±2.7s vs 13.3±1.3s grooming, respectively (p=0.019).

Taken together, our findings show important behavioral effects due to the interactions between light cycle and sex in mice subjected to exercise, compared to sedentary counterparts. Moreover, these results suggest that inverted light cycle conditions might be more suitable for exercise-related studies.

### Baseline corticosterone levels change in a sex and exercise-dependent manner

Corticosterone (CORT) is one of the main stress hormones in rodents, analogous to cortisol in humans^26^. Basal levels of CORT are linked to most physiological processes and strongly modulated by a variety of internal and external cues, including the centrally controlled circadian rhythm^27^. In fact, without stressful stimuli, baseline CORT is released into circulation following a circadian fashion, displaying a significant rise that coincides with the time of awakening^28,29^. Therefore, as rodents are nocturnal animals, the highest levels of daily CORT are usually observed around the time the lights are turned off in most animal facilities that are maintained in a 12/12h cycle with lights ON during daytime^30–32^. CORT is also essential for skeletal muscles to properly function during exercise ^33^. Therefore, we hypothesized that CORT levels may contribute to mediating the observed effects of exercise in different light cycles, as mice kept in REG would be expected to have lower levels of baseline CORT compared to mice kept in INV light conditions. To test this hypothesis, we quantified serum CORT levels at the end of the behavioral evaluation via ELISA (Figure 6A). Our results showed that both exercise and sex had a strong influence on CORT levels (p=0.036 for sex and p=0.003 for exercise, overall Three-way ANOVA), while light cycle did not have a significant effect when considered individually (p=0.634). Notably, the interaction between light cycle and exercise was statistically significant (p<0.0001).

**Figure 6.**
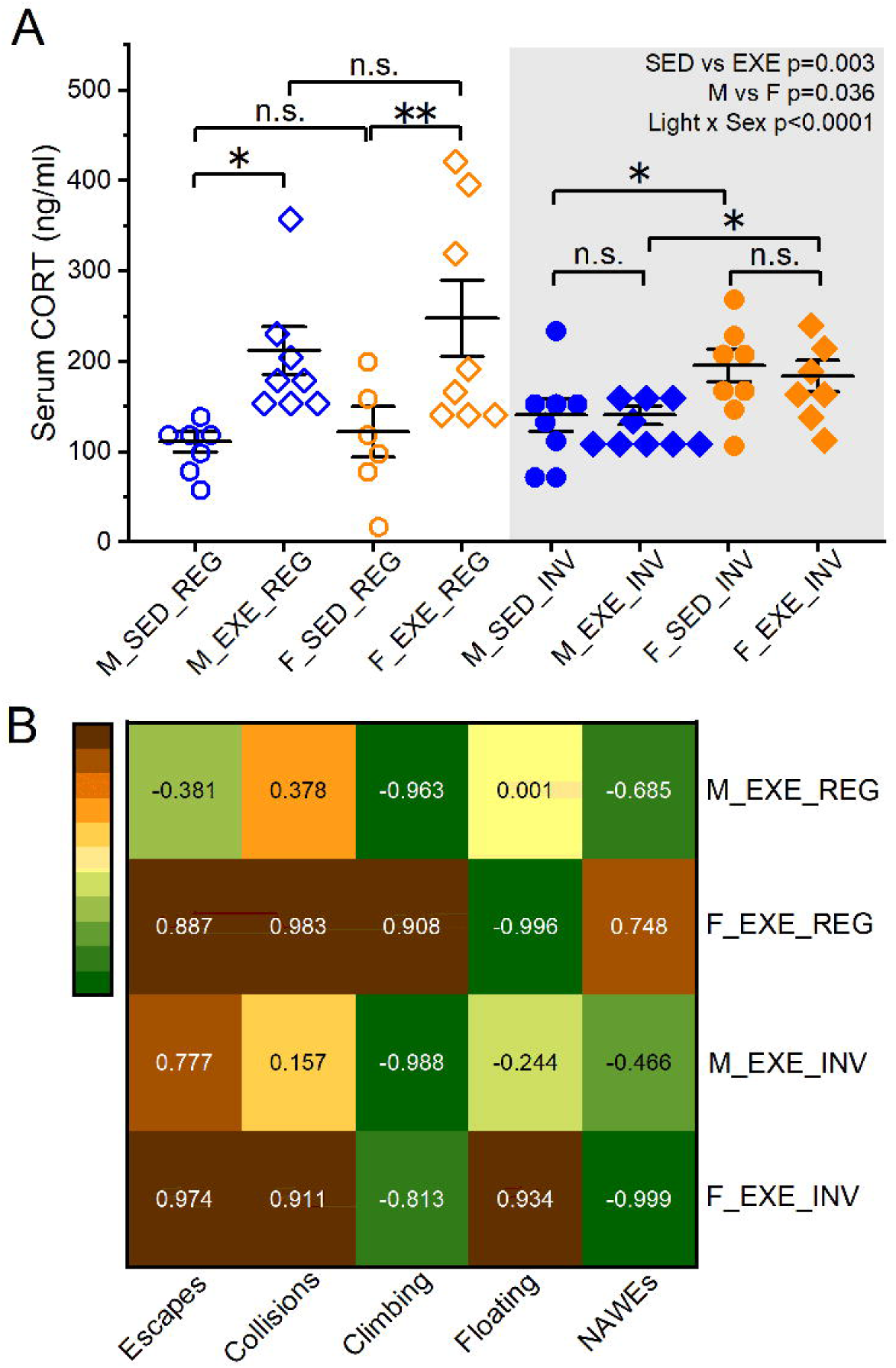
Baseline serum levels of corticosterone (CORT) measured 24 hours after the last day of daily swimming protocol and after the behavioral testing was completed (A). Statistical analysis was conducted by Three-way ANOVA and Tukey’s test post hoc for means comparison. (*) denotes statistical significance with p<0.05 and “n.s.” denotes “not significant” (p>0.05). Panel B shows a heat map graph of Pearson correlation analysis between different behaviors observed during swimming and CORT levels. The numbers in each square represent calculated Pearson coefficient.

Overall, we observed higher baseline CORT levels in mice that exercised and were kept in REG conditions, regardless of sex. However, this presumably exercise-associated increase in CORT was blocked in mice kept in INV light conditions. Furthermore, a sex-dependent difference was revealed as females in the INV light cycle showed higher basal CORT than males, regardless of exercise (p=0.05 between M and F in INV conditions, Three-way ANOVA and Tukey’s Test for means comparison), which was not the case in in mice kept in REG conditions (p=0.74 between M and F). Mice in the M-SED-REG and M-EXE-REG groups had an average serum CORT of 111.3±11.5 vs 212.02±26.5ng/ml (p=0.044, n=7 and 8, respectively). F-SED-REG and F-EXE-REG mice had 121.9±27.9 and 247.6±42.1ng/ml of CORT (p=0.008, n=8 for both groups). On the contrary, mice in the M-SED-INV and M-EXE-INV groups had virtually the same levels of CORT, 140.9±18.2ng/ml vs 140.37±10.65ng/ml respectively (p=0.999, n=8 for both), while F-SED-INV and F-EXE-INV mice had on average 195.8±18.1 vs 183.6±17.2ng/ml, respectively (p=0.999, n=8 and 7 mice respectively).

To visualize the relationship between behaviors displayed during swimming and CORT levels we conducted a Pearson correlation analysis (Figure 6B). Importantly, because we only obtained average values for behavioral scores during swimming (i.e., we manually counted all behaviors in a group per day, not per individual mice), this is an estimation of correlation and not a statistical analysis (and accordingly, we did not calculate statistical significance). Figure 6B shows a heat map representation of the main findings of this correlation comparison. In general, robust sex-dependent differences were observed in some of the active behaviors analyzed, regardless of the light cycle. For example, the number of escapes and collisions was strongly correlated with CORT levels in both REG and INV female groups, not in males. Other behaviors, such as climbing attempts, demonstrated a similar correlation with CORT across both groups of males (REG and INV), but showed a negative correlation in females. Lastly, passive coping behaviors (floating and NAWEs) were modulated not only by sex but also by light cycle, with females showing the most strikingly opposite differences while males showed comparable correlation regardless of the light cycle they were kept in during the experiment.

## DISCUSSION

Here we show that sex and circadian rhythms interact to shape both performance and stereotyped behaviors during swimming exercise in young adult WT mice. Our results extend prior work on time⍰of⍰day–dependent exercise capacity in running paradigms to a different exercise context (i.e., group swimming), linking it with sex⍰specific diurnal modulation of swimming behavior. Furthermore, our findings are novel as most previous studies involving swimming behavior have been conducted in the context of forced⍰swim models of stress to probe “depression-like” behaviors^34^. Unlike these studies, our swimming protocol was designed to minimize stress reactivity while focusing on physical activity. To our knowledge, no previous study has systematically combined light cycle, sex, and detailed analysis of swimming behavior with endocrine and body⍰weight outcomes, making the present work the first to identify a sex⍰ and circadian⍰specific vulnerability of female mice to chronic swimming performed during the rest (REG) period.

Multiple studies have demonstrated that time⍰of⍰day strongly modulates endurance capacity and molecular responses to exercise in mice, with higher performance and distinct transcriptional responses being observed during the early active phase^35,36^. However, most of this prior work in mice has been done using treadmill or wheel running protocols, whereas our paradigm uses repeated swimming in groups. Hence, our exercise paradigm involves distinct postural control, thermoregulatory demands, and social context which are not possible to investigate using running-based exercise paradigms. Thus, our data broaden circadian⍰exercise concepts to another modality of physical activity, allowing us to further study other factors that might influence performance and consequences of exercise. Moreover, our findings that mice kept in REG light conditions that exercised during the same phase (i.e., in their rest phase) showed greater body⍰weight loss and higher CORT responses than mice under INV light conditions (i.e., in their active phase) suggest that implementing swimming exercise during the rest phase enhances physiological cost and stress reactivity.

### Swimming related behaviors are modulated by sex and circadian rhythm

Our study originated from the simple observation that mice kept in different light cycles displayed uniquely distinctive behaviors during repeated swimming exercise. After careful quantification, our analysis revealed important differences in swimming exercise and associated physiological responses. In general, male mice kept in REG light cycle (i.e., lights ON during the day, rest phase) exhibited more coping strategies such as escape attempts, collisions and NAWEs, than male mice kept in INV light (i.e., light OFF during the day, active phase). Conversely, female mice in REG light showed the lowest mobility during swimming compared to females in INV and males in any light cycle, indicating that sex-dependent coping mechanisms are strongly modulated by circadian phase.

Swimming has been extensively used in behavioral research in the context of testing spatial memory (i.e., Morris water maze), or to study responses to inescapable stressors and depression-like behavior (i.e., forced swim test, FST). In addition, swimming is a common model of exercise in rodents, used to probe a variety of physiological outcomes including cardiovascular, skeletomuscular, neurological, etc.^37^. Previous findings have shown that in forced⍰swim paradigms, immobility and active swimming display robust diurnal rhythms with sex⍰dependent characteristics in both WT and clock gene deficient mice^38^, indicating that the circadian system and sex jointly regulate swimming behavior under acute stress. Our data show that a similar interaction is evident during repeated, subacute swimming exercise, where several stereotyped coping behaviors were differentially expressed as a function of sex and light⍰cycle. Interestingly, most behavioral differences were observed under REG light, with relatively attenuated sex effects in mice kept in INV light. This pattern suggests that swimming exercise during the active phase (i.e., INV conditions) may narrow sex differences in behavioral strategies, whereas exercising during the rest phase (REG) unmasks sex⍰specific coping patterns. This aligns with reports that sex and circadian phase jointly modulate daily locomotor rhythms and stress responses in mice^38^, even outside of exercise contexts^39^.

There is currently limited information on mouse swimming behavior outside of the above-mentioned forced stress-inducing paradigms. Moreover, while circadian modulation of behavior in the FST has been reported^40,41^, our study is the first to explicitly compare the effect of the circadian phase on swimming exercise in both male and female mice. In addition, our analysis revealed stereotyped behaviors such as NAWEs that suggest a shift in coping strategy that developed as mice were repeatedly subjected to swimming exercise. Such shift was more prevalent in REG mice which provides another indication that swimming during the rest phase was more stressful than swimming in the active phase (INV). Our observations align with the idea that switching from active to passive coping in the FST might be associated to learning instead of indicating depression-like state and despair^42,24^. Along these lines, a recent report that documented group swimming via video tracking analysis reported that while there was great variability in individual swimming performances, mice that preferred to swim in the central area of the pool had lower CORT levels and showed larger cardiovascular improvements in response to exercise. However, neither the sex nor the specific details of the light cycle used were provided (only the 12h/12h duration)^43^.

### Exercise-associated increase in Corticosterone (CORT) is observed only in mice that exercised during REG phase

Our analysis showed significant differences in baseline CORT levels after swimming protocol. Both male and female mice that were kept in REG conditions and exercised had significantly higher serum CORT levels compared to their sedentary counterparts. However, while exercise did not change CORT levels in mice kept in INV conditions, females had a significantly higher CORT level compared to males regardless of exercise, a major sex-dependent difference that was not observed in mice kept in REG light. These results align with previous literature showing sex differences in the Hypothalamic-Pituitary-Adrenal (HPA) axis responsiveness and glucocorticoid secretion. Several studies have shown that female mice have higher CORT levels compared to males due to a combination of factors including greater intrinsic HPA axis sensitivity, hormonal and ovary-derived factors that influence stress responsiveness, and sex differences in glucocorticoid receptor feedback sensitivity^44,45^.

CORT release follows a well-known circadian fluctuation in most mammals, which is tightly regulated by the HPA axis (reviewed in^46^ for rodents and^47^ for humans). During the early light/rest phase (ZT0-ZT6), when REG mice swam in our study, CORT levels are typically low. Conversely, right before the beginning of the early dark/active phase (ZT12-ZT18), when INV mice swam, CORT levels peak and reach their highest daily levels^48^. Consistent with this established pattern, we expected SED mice to exhibit lower CORT levels in REG light and higher in INV. However, while there was a trend, our results showed no statistically significant differences. This discrepancy might be related to the time of blood collection in the dark phase, which was 2-3h after CORT levels peaked (and thus CORT levels were already decreasing). Remarkably, the fact that swimming exercise increased CORT only in mice kept in REG could still be related to the known circadian modulation of CORT release^29^. In other words, mice kept in REG conditions that exercised showed larger increase in CORT than mice kept in INV light as CORT production is likely higher at the beginning of the dark/active period in rodents (i.e., INV light in our study) and thus this lack of exercise-induced increase could indicate a ceiling effect in CORT release. This idea is supported by other studies showing a larger increase in CORT in response to different stressors during the beginning of the light or rest phase^48–50^. Similarly, Per2 knockout mice that have a dysregulated circadian rhythm, also display increased CORT levels and depressive-like behaviors^51^. Taken together, our data suggests that swimming performed during the light/rest phase (REG) can act as an acute physiological stressor, leading to cumulative higher CORT levels. Considering that this increase in CORT in exercised mice was not observed in mice kept in INV light (dark/active phase), these findings collectively highlight the strong circadian and sex-dependent modulation of stress reactivity.

### Circadian modulation of body weight changes in response to swimming exercise

We observed significant differences in body weight (BW) in response to swimming exercise. Females kept in REG light were the only group that lost weight after 3 weeks of daily swimming, even though this group moved less during swimming sessions. This group also had the highest levels of CORT after chronic swimming, compared to sedentary matched controls. Elevated CORT levels have been observed in models of chronic swimming and other exercise paradigms and have been associated with lower weight gain or reduced adiposity and altered HPA axis function in mice ^52^, particularly in metabolically challenged models^53,54^. Likewise, chronic or repeated stress protocols that elevate CORT typically attenuate BW gain or induce weight loss ^55,56^. Conversely, female mice kept in INV showed no significant change in BW after exercise. Moreover, male mice kept in different light cycles showed opposite effects on BW, namely, males in REG cycle that exercised gained less weight than sedentary controls while males in INV cycle gained more weight that sedentary groups. These findings are consistent with previous work showing that exercise not always leads to weight loss, particularly in younger individuals^57^. Furthermore, these striking differences in BW in response to exercise add another layer of complexity to the interaction between sex, circadian phase and exercise.

Interestingly, we also observed distinctive BW changes between control sedentary (SED) groups kept in different light cycles. Specifically, both male and female sedentary mice kept in REG light gained significantly more weight than mice kept in INV light. Many studies in both animal models and humans have shown that circadian disruptions lead to detrimental metabolic and endocrine alterations. For instance, nightshift workers have a higher rate of cardiovascular diseases, diabetes, obesity, and cancer^58–60^, Similarly, transgenic mice lacking the *clock* gene have a disrupted light dark cycle, are hyperphagic and develop obesity and metabolic syndrome^61^. Our results are aligned with these previous findings because while sedentary mice were not subjected to swimming exercise, their sleep was perturbed daily by handling during BW monitoring. Such changes in sleep patterns are comparable with models of sleep fragmentation, which lead to increased food intake and obesity in mice^62^. In addition, this daily routine was likely perceived as a form of mild chronic stress paradigm, which can also lead to metabolic alterations and weight gain^63^, contrasting with severe chronic stress paradigms that generally cause weight loss in mice^64^.

Taken together, our findings indicate important metabolic adaptations to chronic exercise (or lack of thereof), that are strongly modulated by circadian rhythms and sex, and the interaction between all these factors.

### Females kept in INV cycle showed altered exploratory behaviors in the open field test (OFT)

Notably, behaviors in the OFT were most altered in mice kept in INV light conditions, particularly females, indicating that swimming during active phase can produce robust changes in exploratory/locomotor behavior despite a lower apparent stress load (i.e., no change in CORT or weight after exercise in INV females). However, these exercised INV females had higher CORT levels compared to INV males and sedentary REG males. This dissociation underscores that the behavioral consequences of timed exercise are not fully captured by classical stress readouts (such as CORT levels) and may depend on sex⍰specific adaptations.

In contrast, females that exercised during the REG light cycle displayed increased anxiety-like behavior, as evidenced by reduced time spent in the center of OFT. Although exercise is frequently associated with anxiolytic effects, this indicates that its impact is time-of-day dependent^65^. Growing evidence indicates that circadian disruption can potentiate stress and anxiety-related behaviors through dysregulation of stress-responsive systems, including the HPA axis^66^. Thus, exercise performed during the rest phase (i.e., REG conditions) may act as an additional physiological stressor (likely to cause further circadian dysregulation) rather than a protective factor. However, these effects are also modulated by sex as male mice tested during REG light cycle also exhibited elevated CORT levels, without the anxiety-like phenotype observed in females. This discrepancy between endocrine and behavioral response is consistent with the evidence that glucocorticoid concentration does not linearly predict anxiety-like related outcomes and that this effect depends on other factors such as sex and emotional state^67^. Other studies have shown that females exhibit heightened HPA axis reactivity and altered glucocorticoid feedback regulation, which may render them more vulnerable to the behavioral consequences of external cues such as circadian disruptions^44,45,68^. Thus, exercise performed during the rest phase (REG) may act as a physiological stressor that disproportionately affects females, even when hormonal elevations are observed in both sexes.

Lastly, other exploratory behaviors such as rearing and grooming also showed changes in response to exercise that largely reflected the interaction sex and circadian rhythms. It is notable that SED females in the INV light cycle had increased grooming time compared to REG sedentary females, and that swimming exercise reversed that change. Excessive grooming is typically a behavioral phenotype associated to the stress response in rodents^69,70^, therefore, this change could indicate anxiolytic effects of exercise that were not observed in mice kept and exercised in REG cycle. Altogether, our data further supports the idea that exercise during the rest phase acts as an additional stressor.

### Limitations and Conclusions

There are several methodological considerations in our study. First, swimming behavior was scored manually instead of with video⍰tracking software, which could introduce human error or observer bias. While we used consistent criteria and multiple observers to minimize this limitation, future studies should use blind, automated analysis based on video analysis to improve reproducibility. Second, future studies should also consider monitoring the expression of clock genes in mice exercising at different times of the day, to properly reference circadian involvement. Despite these constraints, our data (together with existing evidence that exercise timing modulates muscular, adipose, and systemic responses) supports the notion that the circadian phase is a determinant factor for exercise as a potential beneficial lifestyle intervention. To account for this, preclinical exercise studies should explicitly control and report circadian phase. Furthermore, our results highlight not only the importance of considering experimental conditions for a variety of behavioral studies, but also how environmental cues have a powerful influence on the outcome of these studies.

